# SARS-CoV-2 Nsp16 activation mechanism and a cryptic pocket with pan-coronavirus antiviral potential

**DOI:** 10.1101/2020.12.10.420109

**Authors:** Neha Vithani, Michael D. Ward, Maxwell I. Zimmerman, Borna Novak, Jonathan H. Borowsky, Sukrit Singh, Gregory R. Bowman

**Affiliations:** Department of Biochemistry and Molecular Biophysics, Washington University School of Medicine, St. Louis, Missouri 63110, United States; Center for Science and Engineering of Living Systems (CSELS), Washington University in St. Louis, St. Louis, Missouri 63130, United States; Medical Scientist Training Program, Washington University in St. Louis School of Medicine, St. Louis, Missouri 63110, United States

**Author notes:** These authors contributed equally to this work (Neha Vithani, Michael D. Ward).

## Abstract

Coronaviruses have caused multiple epidemics in the past two decades, in addition to the current COVID-19 pandemic that is severely damaging global health and the economy. Coronaviruses employ between twenty and thirty proteins to carry out their viral replication cycle including infection, immune evasion, and replication. Among these, nonstructural protein 16 (Nsp16), a 2’-O-methyltransferase, plays an essential role in immune evasion. Nsp16 achieves this by mimicking its human homolog, CMTr1, which methylates mRNA to enhance translation efficiency and distinguish self from other. Unlike human CMTr1, Nsp16 requires a binding partner, Nsp10, to activate its enzymatic activity. The requirement of this binding partner presents two questions that we investigate in this manuscript. First, how does Nsp10 activate Nsp16? While experimentally-derived structures of the active Nsp16/Nsp10 complex exist, structures of inactive, monomeric Nsp16 have yet to be solved. Therefore, it is unclear how Nsp10 activates Nsp16. Using over one millisecond of molecular dynamics simulations of both Nsp16 and its complex with Nsp10, we investigate how the presence of Nsp10 shifts Nsp16’s conformational ensemble in order to activate it. Second, guided by this activation mechanism and Markov state models (MSMs), we investigate if Nsp16 adopts inactive structures with cryptic pockets that, if targeted with a small molecule, could inhibit Nsp16 by stabilizing its inactive state. After identifying such a pocket in SARS-CoV-2 Nsp16, we show that this cryptic pocket also opens in SARS-CoV-1 and MERS, but not in human CMTr1. Therefore, it may be possible to develop pan-coronavirus antivirals that target this cryptic pocket.

**Statement of Significance:** Coronaviruses are a major threat to human health. These viruses employ molecular machines, called proteins, to infect host cells and replicate. Characterizing the structure and dynamics of these proteins could provide a basis for designing small molecule antivirals. In this work, we use computer simulations to understand the moving parts of an essential SARS-CoV-2 protein, understand how a binding partner turns it on and off, and identify a novel pocket that antivirals could target to shut this protein off. The pocket is also present in other coronaviruses but not in the related human protein, so it could be a valuable target for pan-coronavirus antivirals.

## Introduction

With the coronavirus 2019 (COVID-19) pandemic ravaging communities across the globe there is a massive ongoing effort to understand the molecular machinery of coronaviruses, which may provide insight into therapeutic opportunities (1–3). The severe acute respiratory syndrome coronavirus 2 (SARS-CoV2) virus responsible for COVID-19 disease has infected over sixty million and killed over 1.5 million people globally to date (4). Additionally, coronaviruses have caused several past epidemics including severe acute respiratory syndrome (SARS) and Middle East respiratory syndrome (MERS) which had fatality rates of ∼10% and ∼34%, respectively (5, 6). Therefore, there is likely to be evolution and outbreaks of additional zoonotic coronaviruses in the future (7). While vaccine trials for COVID-19 are successfully wrapping up, there are still no approved antivirals that reduce mortality to coronavirus infections (8–10). Taken together, there is strong incentive to understand the fundamental mechanisms of how these coronaviruses operate in hopes of discovering effective therapeutics. Biophysical studies can provide these details, and a tremendous amount of biophysical work has already been done to understand the virus’ twenty-nine proteins. So far, the spike protein, positioned on the outside of the viral envelope, has proven to be a good vaccine candidate (11). Beyond the spike, the sixteen “nonstructural” (i.e. accessory) proteins carry out the majority of the virus’ essential processes, making them good targets for antiviral therapeutics (12, 13).

Among the nonstructural proteins (Nsp’s), Nsp16 is particularly important to the viral replication cycle as it is essential to coronavirus’ immune evasion (14–16). Nsp16 is a 2’-O-Methyltransferase (2′-O-MTase) that forms part of the replication-transcription complex (17). It mimics the human protein Cap-specific mRNA (nucleoside-2’-O-)-methyltransferase (CMTr1) to perform a crucial step in capping transcribed mRNA (18). Specifically, Nsp16 facilitates the transfer of a methyl group from its S-adenosylmethionine (SAM) cofactor to the 2’ hydroxyl of ribose sugar of viral mRNA (18, 19). This methylation both improves translation efficiency and camouflages the mRNA so that it is not recognized by intracellular pathogen recognition receptors, such as IFIT and RIG-I (15, 20). Importantly, inhibiting or knocking out 2′-O-MTase activity severely attenuates viral replication and infectivity of coronaviruses (13, 20). Thus, developing small molecules inhibitors of Nsp16 is a promising therapeutic strategy.

Interestingly, while all other 2′-O-MTases (eukaryotic and viral) are active as monomers, Nsp16 requires a binding partner, Nsp10, to be active (16–18, 21–23). In fact, Nsp16 does not even bind its ligands (SAM and RNA) in the absence of Nsp10. In the experimentally-derived structures of the Nsp16/Nsp10 complex, Nsp10 does not form any direct interaction with either ligand (Fig. 1a), suggesting that Nsp10 may allosterically regulate Nsp16 to enable substrate binding (18, 19, 24–27). Given that there is significant structural variation in the RNA-binding loops of different crystal structures of Nsp16 (Fig. 1b) and structures of monomeric Nsp16 have not been solved, we hypothesized that Nsp16 is highly dynamic in solution, and Nsp10 acts by stabilizing the active state. In contrast, we anticipate that human CMTr1 would be less dynamic as it doesn’t require a binding partner for substrate binding and has been crystalized in its monomeric state. Often, dynamics of proteins reveal allosteric pockets that remain hidden in their crystal structures (i.e., cryptic pockets). If monomeric Nsp16 is more dynamic than CMTr1, it may adopt inactive configurations that reveal allosteric cryptic pockets, which can be targeted by small-molecule inhibitors for its selective inhibition.

**Figure 1.**
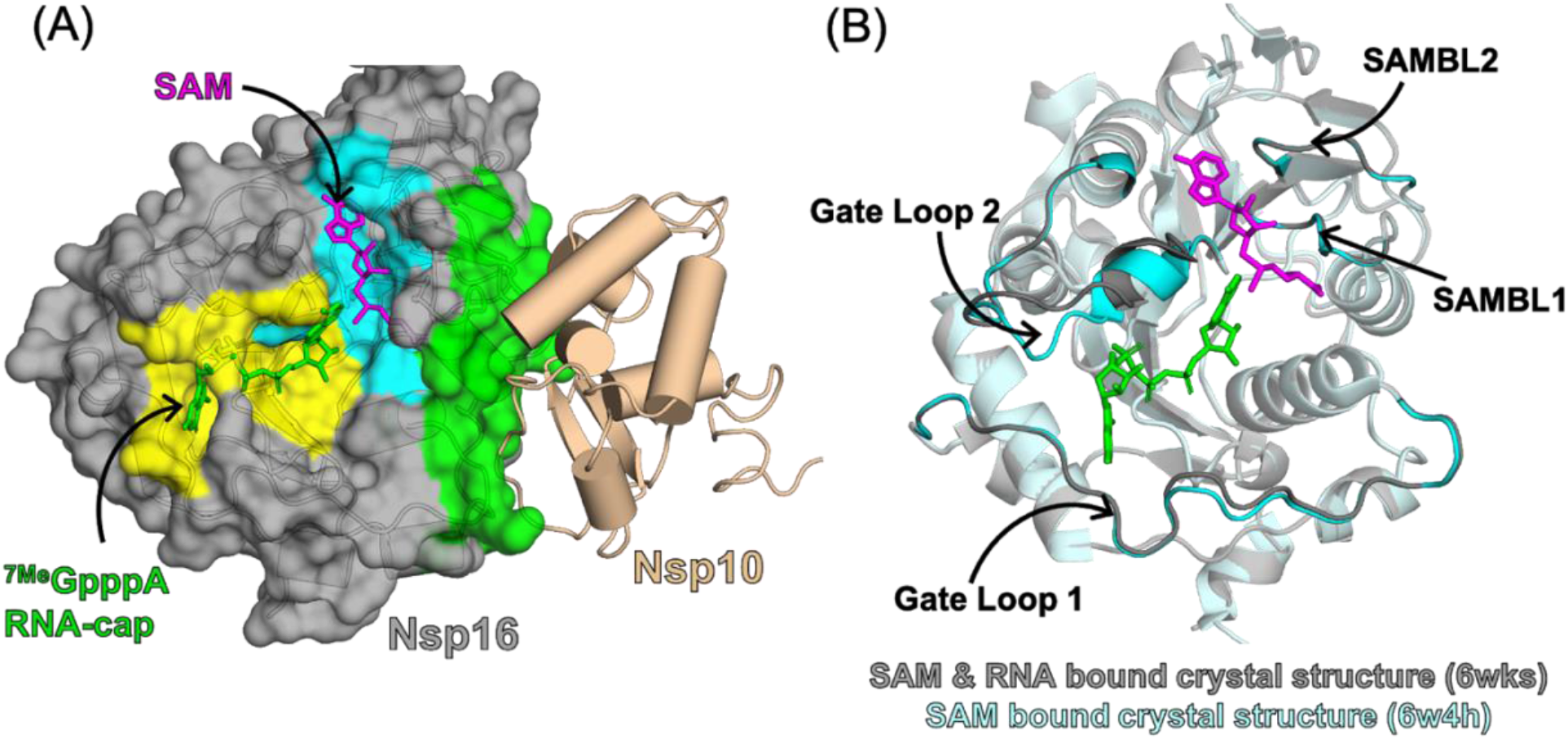
Substrate binding pockets and Nsp10 binding interface of Nsp16 observed in the crystal structure of the Nsp16/Nsp10 complex (PDB: 6wks). (A) Surface representation of Nsp16 showing the SAM-binding pocket (cyan), RNA-binding pocket (yellow) and Nsp10-binding interface (green). (B) Overlay of Nsp16 structures from structures of the Nsp16/Nsp10 complex with RNA (PDB: 6wks, shown in grey) and without RNA (PDB: 6w4h, shown in cyan), showing structural heterogeneity in the RNA-binding site. Gate loop 1 and Gate loop 2 of the RNA-binding pocket, and SAM-binding loop 1 (SAMBL1) and SAM-binding loop 2 (SAMBL2) lining the SAM-binding pocket are highlighted.

Here, we use computer simulations to understand the activation mechanism of Nsp16 and identify cryptic pockets that may be valuable antiviral targets. Active site inhibitors, such as Sinefungin, have been shown to outcompete SAM binding and render Nsp16 catalytically inactive (28, 29). However, there are more than 200 human proteins with known or putative methyltransferase activity that use SAM as a cofactor (30). Therefore, it may be difficult to design antivirals that target the SAM (or RNA) binding sites of Nsp16 without eliciting off-target effects by also binding human methyltransferases. For example, Sinefungin has been shown to occupy the SAM-binding pocket of human N7 methyltransferase in a crystal structure (PDB: 3epp). Targeting the Nsp16/Nsp10 interface could be an alternative means to selectively inhibit Nsp16 since CMTr1 lacks a homologous binding partner. Towards this, peptide-based inhibitors that mimic Nsp10 to compete for interactions at the Nsp10/Nsp16 interface have been shown to inhibit Nsp16 activity (31, 32). While this approach seems promising, peptide-based inhibitors face challenges including limited stability and shelf-life, the possibility of adverse immunogenic responses, and the high cost of production (33). To expand the therapeutic opportunities, we search for other ways to inactivate Nsp16. First, we compare the structure and dynamics of SARS-CoV2 Nsp16 in the presence and absence of Nsp10 to understand Nsp16’s activation. Specifically, we use over one millisecond of molecular dynamics simulation data (2) to characterize how Nsp10 binding shifts Nsp16’s conformational ensemble to activate Nsp16. After showing that the resulting model is consistent with a variety of experimental observations, we use it to hunt for cryptic pockets that may provide a means to inhibit Nsp16. Finally, we extend our simulations to SARS-CoV-1, MERS, and human CMTr1 to determine if targeting such a pocket could provide an opportunity to develop pan-coronavirus antivirals.

## Results and Discussion

### Nsp10 promotes opening of Nsp16’s SAM- and RNA-binding pockets

While experimental studies have demonstrated that Nsp16 requires Nsp10 to be functionally active, the structural determinants of Nsp16’s activation remain unknown (17, 18, 23). Chen et. al. proposed that Nsp10’s stimulatory effects are rooted in its ability to assist Nsp16 in binding SAM and RNA, which is supported by data showing that Nsp16 alone cannot bind SAM or RNA (18). They also propose that Nsp10 manages this by stabilizing or changing the conformation of the SAM binding pocket based on the fact that Nsp10 contacts SAM binding loops in their crystal structure (and numerous other structures). However, without assessing Nsp10-Nsp16 complex’s dynamics and comparing it to monomeric Nsp16, this hypothesis is left wanting. It has also been proposed that Nsp10 assists in RNA binding by directly contacting RNA (34). However, a recent crystal structure with RNA bound (PDB: 7jyy) contains a stretch of nucleotides long enough to contact Nsp10, but the RNA curls off into solution instead of interacting with Nsp10. Another recent study compared an RNA and SAM bound Nsp10/16 complex structure to one with only SAM bound and found a major opening of RNA binding gate loops suggesting that the dynamics of these loops might be important for Nsp16 activation (25). However, it is not clear if Nsp10 plays a role in those dynamics. Altogether, there is strong evidence that Nsp10 modulates Nsp16’s structure and dynamics to assist it in binding SAM and RNA, but the mechanism of these structural changes is unclear.

To explore how Nsp10 activates Nsp16, we analyzed simulations of Nsp16 in the presence and absence of Nsp10 using DiffNets. Recently, our group combined the sampling powers of the FAST-pockets adaptive sampling algorithm (35) and the computational resources of Folding@home to accumulate more than one millisecond of simulation data between simulations of monomeric Nsp16 and the Nsp16/Nsp10 complex (see methods) (2). Here, we compare these simulations using a deep learning-based dimensionality reduction algorithm called DiffNets (36). DiffNets has been shown to accurately capture the structural determinants of biochemical differences between protein variants. While we are not considering protein variants, our problem is similar since Nsp16 has different biochemical properties when in the presence/absence of Nsp10 (i.e. active/inactive). Therefore, we trained a DiffNet to learn the structural determinants of Nsp16 activation by learning differences between Nsp16’s ensemble when in the presence and absence of Nsp10. For each simulation frame, the DiffNet learns a low dimensional projection of the protein structure and classifies the structure with a label between 0 and 1 that indicates the likelihood that the structure is associated with Nsp16 being active.

Analysis of the DiffNet suggests that Nsp10 shifts Nsp16’s conformational ensemble to stabilize more open SAM- and RNA-binding pockets. Using the DiffNet classification labels, we identified ten structures that are representative of the progression from Nsp16 inactive states to active states (see methods and Fig. 2). We noticed that RNA gate loop 2 moves away from RNA gate loop 1, making for a more open RNA binding pocket in active states compared to inactive states (Fig. 2A). Additionally, the SAM-binding pocket also opens up in the active states relative to the inactive states. RNA-binding gate loop 2 and SAM-binding loop 2 move away from each other in the active state, which widens the pocket creating space for SAM. (Fig. 2A). Strikingly, the structure associated with the highest label (i.e. most strongly associated with Nsp16 activation) matches well to a recently solved crystal structure that is bound to both RNA and SAM (Fig. 2B) (25). This DiffNet result is encouraging since the simulations were started from a markedly different crystal structure (no RNA bound) and had no a priori information about the RNA bound structure. This result implies that the DiffNet learned that Nsp10 activates Nsp16, in part, by rearranging the RNA gate loop into an RNA binding competent pose. Though it is known that this RNA gate loop needs to open to bind RNA, this is the first evidence, to our knowledge, to suggest that Nsp10 may activate Nsp16 through increasing its propensity to form a more open RNA-binding pocket. Altogether, these results suggest that Nsp10’s presence increases the propensity for both SAM- and RNA-binding pockets to be open.

**Figure 2.**
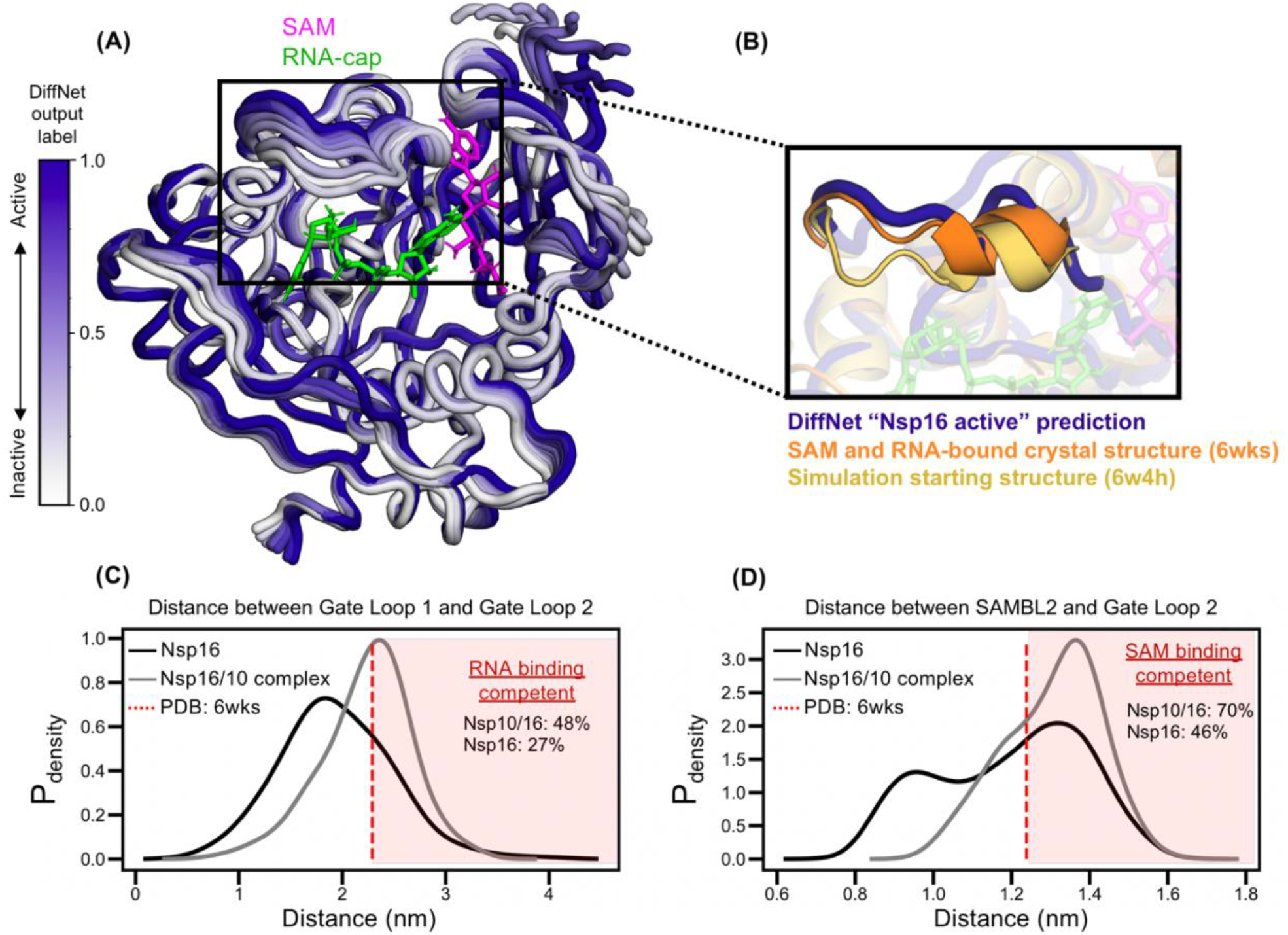
Nsp10 binding shifts Nsp16’s conformational ensemble increasing its propensity to adopt structural states that are ligand binding compatible. (A) Ten structures of Nsp16 that represent the DiffNet prediction changing from inactive to active (white to purple). (B) Comparison of the DiffNet predicted active state (purple) to the starting simulation state (yellow) and a known RNA bound structural state (orange). (C) Probability-weighted distance distribution between RNA-binding gate loops 1 and 2 comparing monomeric Nsp16 (black) to the Nsp10-Nsp16 complex (gray). (D) Probability-weighted distance distribution between SAM-binding loop 2 and gate loop 2, comparing monomeric Nsp16 (black) to the Nsp10-Nsp16 complex (gray). For (C) and (D), the distance for a SAM and RNA bound crystal structure is also plotted (red dotted line).

To quantify the effect of Nsp10 on the SAM- and RNA-binding pockets, we built MSMs for both the complex and monomeric Nsp16. MSMs are a statistical framework for analyzing molecular dynamics simulation data that provide (among other things) a discrete map of structural configurations, an equilibrium population value that corresponds to the proportion of time a protein spends in any given configuration, and the probability of transitioning between any pair of configurations (37). We constructed MSMs for Nsp16 simulations both in the presence and absence of Nsp10.

Our MSMs reveal that Nsp10 binding stabilizes open structures of both the SAM- and RNA-binding pockets that are competent to bind their respective substrates. We first found that the presence of Nsp10 results in a substantial reduction of flexibility in important binding components including both SAM binding loops and RNA gate loops (see SI, Fig. S1). This result is somewhat surprising since gate loop 2, which contacts both SAM and RNA, is not in direct contact with Nsp10, suggesting strong allosteric communication. Next, we calculated the distribution of distances for opening and closing of the SAM and RNA binding pockets (Fig. 2C,D). From these histograms it is clear that both of these binding pockets have an increased propensity to open when Nsp10 is present. We considered pockets as SAM/RNA binding competent when the distance between loops in a pocket is at least as open as in the crystal structure that binds both ligands (PDB: 6wks). From this analysis, Nsp16 adopts binding competent states with higher probability when Nsp10 is present vs when Nsp10 is absent for both SAM (0.70 ± 0.04 vs 0.46 ± 0.04) and RNA (0.48 ± 0.04 vs 0.27 ± 0.03). Altogether, our data suggest that Nsp10 aids SAM and RNA binding by preventing the collapse of SAM and RNA binding gate loops. Our analysis also provides structural snapshots of what inactive states look like, which may be useful in targeting Nsp16 with therapeutics.

### A cryptic pocket in Nsp16 is a potential therapeutic target

A traditional approach to drug development involves molecules designed to target binding cavities observed in singular structural snapshots of a protein, but this approach often misses “cryptic” pockets that can form in proteins due to thermal fluctuations. Often times the active site of an enzyme is targeted for drug development to design an inhibitor that can outcompete substrate binding. However, active sites are often conserved among functional homologs. In the case of Nsp16, its human homolog (CMTr1) shares the same overall fold and binds the same substrates. Though there are significant sequence and structural differences in the active site, specificity may be more easily achieved by targeting a less functionally relevant region of the protein. Cryptic pockets can provide both a new target for drug development and the potential to achieve specificity. For example, cryptic pockets that remain closed and invisible in the crystal structure, but open in solution due to thermal fluctuations (38), can present unique potential binding sites due to differences in the dynamics of subsets of homologs (e.g. open in coronavirus homologs, but closed in human CMTr1). Therefore, it may be easier to achieve specificity by targeting a cryptic pocket. Importantly, the cryptic pocket must communicate with functional sites in order for it to be an effective therapeutic target. Here, we explore if Nsp16 contains any cryptic pockets that, when open, would stabilize the inactive state identified with DiffNets.

To find cryptic pockets, we applied “Exposons”, an algorithm (38) that identifies residues with cooperative changes in solvent exposure, to Nsp16 simulation data. Using this method, we found that residues in the β3 strand and α3 helix transition between closed states and open states (i.e. low to high solvent accessible surface area) (Figure 3A). Specifically, the β4 strand curls up to form an α-helical structure, which results in surface exposure of β3 and residues from α3 (Fig. 3A). The opening motion of β4 shifts the adjacent SAMBL2 against gate loop 2 to collapse the SAM binding pocket in a closed conformation (Fig. 3A,B). This agrees with the DiffNet prediction that the β4 strand moving away from β3 is associated with inactivation (see SI. Fig. S2). Further, several residues forming this cryptic pocket directly contact Nsp10 in crystal structures of the Nsp16/Nsp10 complex (see SI. Fig. S3). The β3-β4 pocket opening displaces these Nsp10 binding residues, which could inhibit Nsp16’s association with Nsp10 (see SI. Fig. S3). Finally, we find that this open pocket structure is commonly visited as part of monomeric Nsp16’s conformational ensemble, as measured with MSM equilibrium populations (Fig. 3B). Taken together, we propose that targeting the β3-β4 pocket with a small molecule could inhibit Nsp16’s activity by preventing SAM binding or preventing association with Nsp10.

**Figure 3.**
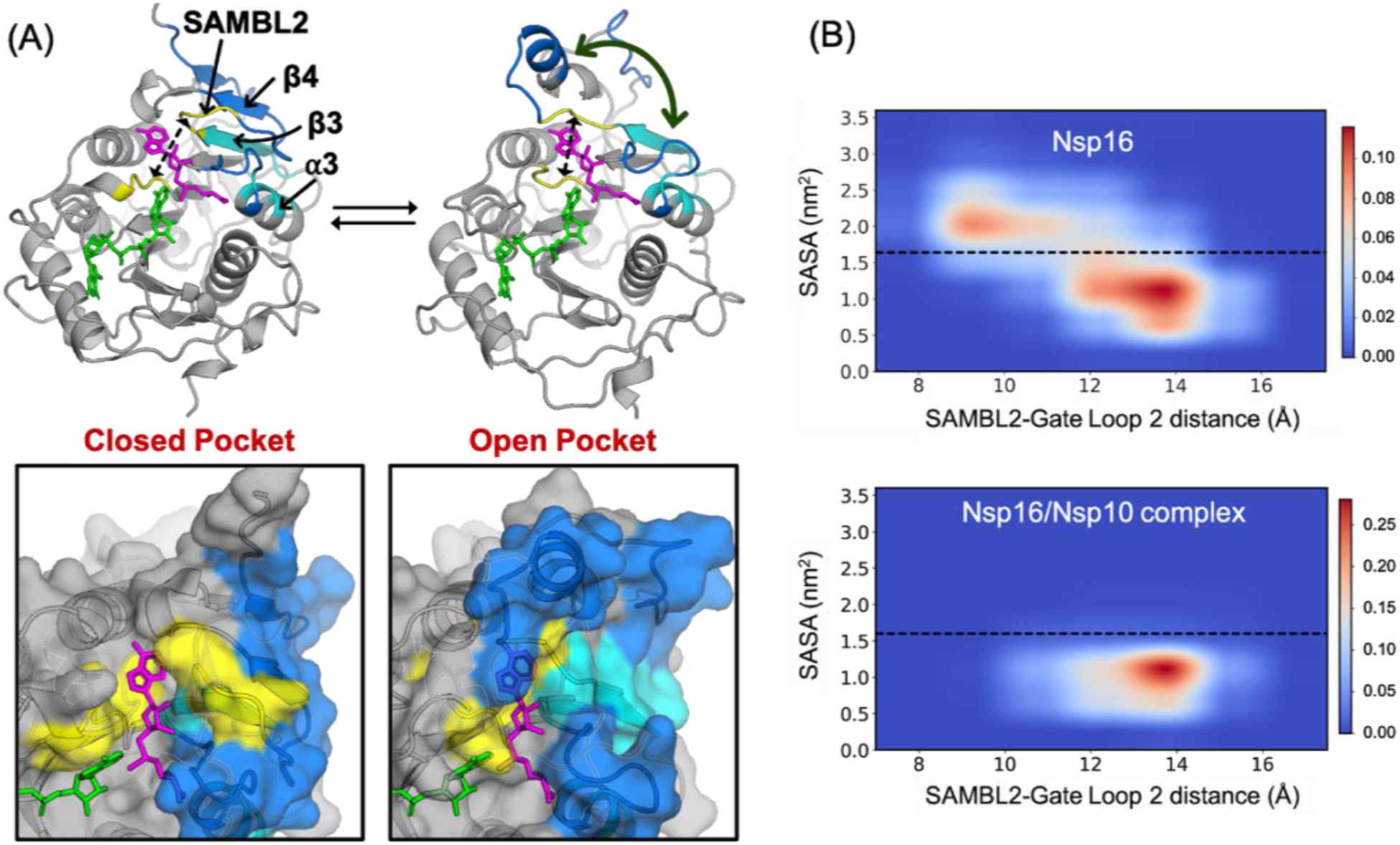
Cryptic pocket opening in SARS-CoV-2 Nsp16. (A) Structural states with the cryptic pocket closed and open. The insets show surface views of the closed and open pocket. Residues exposed upon pocket opening are shown in cyan and the regions undergoing the opening motion are shown in blue. Collapse of the SAM-binding pocket is measured as the distance between SAMBL2 and gate loop 2, shown in yellow. (B) Equilibrium probability weighted 2D histograms of solvent-accessible surface area (SASA) of pocket residues (shown in cyan in A) and the distance between SAMBL2 and gate loop 2 in Nsp16 for monomeric Nsp16 (upper panel) and the Nsp16/Nsp10 complex (lower panel). The black dotted line separates the pocket closed and open states in Nsp16.

### Conservation of the cryptic pocket in Nsp16 makes it a promising target for broad-spectrum inhibitors

To explore the possibility of targeting the cryptic pocket for broad-spectrum inhibition of coronaviruses, we evaluated the conservation of cryptic pocket opening in Nsp16 homologs. Ideally, a therapeutic developed to treat SARS-CoV-2 would also work against other coronaviruses like MERS, SARS-CoV-1, and potentially future outbreaks. Additionally, the therapeutic target should be sufficiently dissimilar from human CMTr1 such that it would not cause unwanted, off-target effects. While we identified a promising cryptic pocket in SARS-CoV-2, we wanted to investigate if this pocket is specific to SARS-CoV-2, or specific to coronaviruses in general, or if it is common across homologs including CMTr1.

First, we analyze cryptic pocket conservation by comparing sequence features and structural features based on the native, folded state. We find that the β3-β4 pocket residues are 100% conserved between SARS-CoV-2 and SARS-CoV-1 (Fig. 4B). Additionally, of the eleven residues that form the pocket, there are only two non-conservative mutations between SARS-CoV-2 and MERS. Based on the sequence similarity, we expect that, if the cryptic pocket forms in all homologs, it may be possible to develop small-molecule therapeutics that targets all three. Further, we find substantial sequence differences between SARS-CoV-2 and CMTr1. Eight out of the eleven pocket residues are non-conservative mutations relative to SARS-CoV-2. Based on sequence differences alone, we reason that selective inhibition could be achieved even if the cryptic pocket is adopted by CMTr1. Moreover, the sequences and structure of SARS-CoV-2 Nsp16 and human CMTr1 are sufficiently different in the β3-β4 pocket region that the human protein may not even have the cryptic pocket (SI, Fig. S4). Based on these sequence and structural differences, combined with the lack of requirement of a stabilizing binding partner, we hypothesized that cryptic pocket opening is not likely to be conserved in CMTr1.

**Figure 4.**
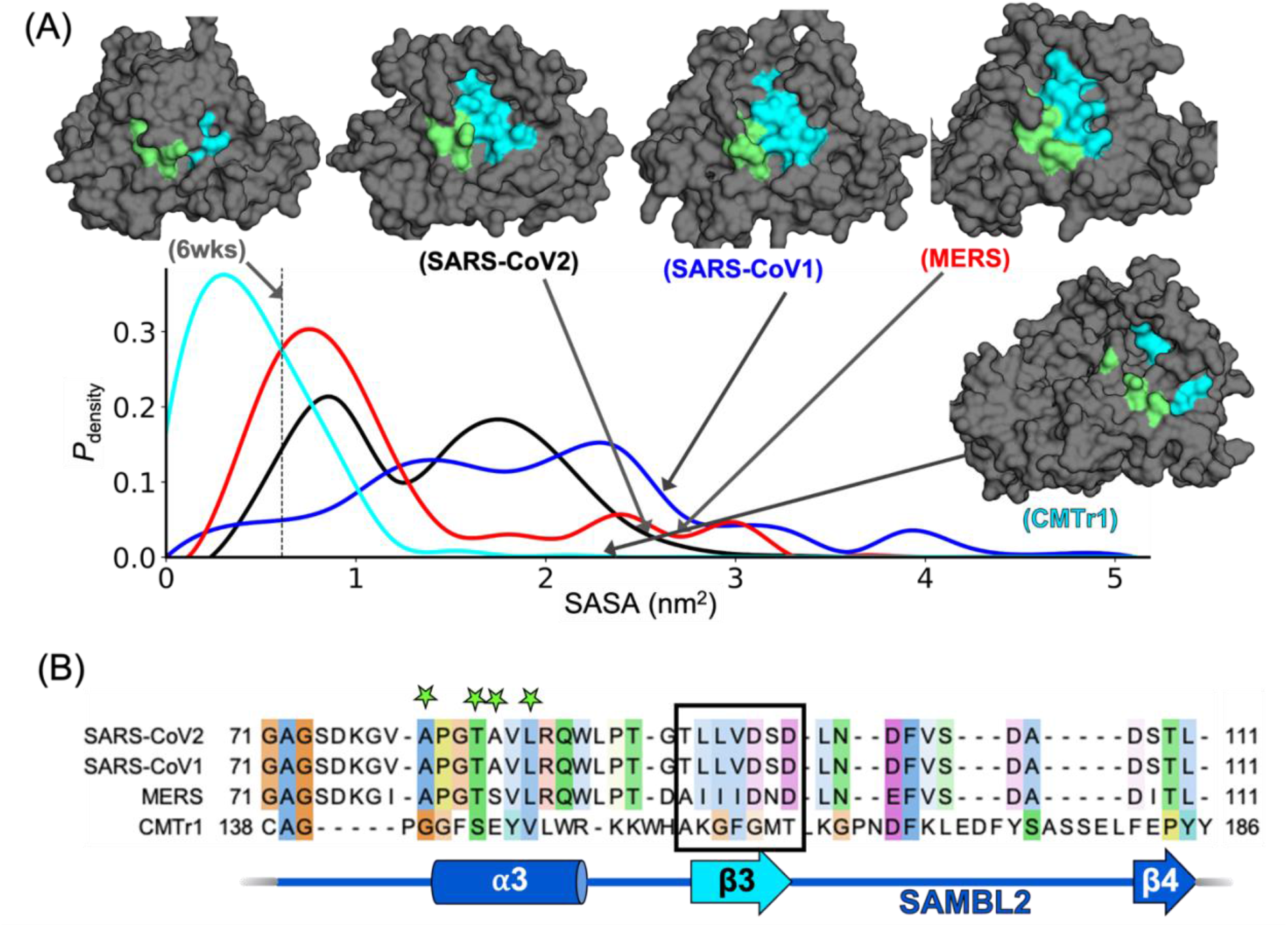
Comparison of cryptic pocket opening in Nsp16 homologs and human CMTr1. (A) Equilibrium probability-weighted distribution of the solvent exposure of pocket forming residues for SARS-CoV2 (black), SARS-CoV1 (blue), MERS (red) and CMTr1 (cyan). Structures representing the open pocket are shown for each homolog with β3 colored in cyan, and other pocket forming residues from alpha3 colored in green. Black dotted line depicts SASA of pocket residues in the crystal structure of Nsp16/Nsp10 complex (PDB: 6wks). (B) Structure-based sequence alignment of Nsp16 homologs (SARS-CoV2, SARS-CoV1 and MERS) and human CMTr1 is shown for the cryptic pocket forming regions. Residues of Beta3 are are marked inside the black colored box, and other pocket forming residues from alpha3 are by green colored stars.

To explore cryptic pocket opening across homologs, we performed FAST-pocket simulations of monomeric Nsp16 for SARS-CoV1 and MERS, as well as, for human CMTr1. Then, we built an MSM for each homolog and measured opening of the β3-β4 pocket by measuring the equilibrium weighted solvent exposure of the pocket residues we previously used to define the pocket (i.e. Fig. 3). In these simulations, we find that the β3-β4 pocket opens with high probability in both SARS-CoV1 and MERS Nsp16 (Fig. 4). Interestingly, the pocket is most likely to open in SARS-CoV-1, followed by SARS-CoV-2, then MERS. Encouragingly, we find that the β3-β4 pocket has a substantially lower probability of opening in CMTr1. Taken together, features of the β3-β4 cryptic pocket in coronavirus homologs of Nsp16 appear sufficiently similar to each other and dissimilar to CMTr1 to make for a promising target for pan-coronavirus inhibitors.

## Conclusions

Our work provides mechanistic insight into how Nsp16 is activated and reveals a new opportunity for inhibiting this essential viral component that could provide a target for pan-coronavirus antivirals. First, we elucidate the activation mechanism of Nsp16 by comparing its dynamics in the presence and absence of its activator, Nsp10. Our results are consistent with previous experimental findings that Nsp16 cannot bind its substrates SAM or RNA in the absence of Nsp10 (18). We provide a structural rationale for this observation by elucidating the structural dynamics of Nsp16 in its monomeric state, which has remained inaccessible to experimental studies, and comparing it to the structural dynamics of the Nsp16/Nsp10 complex. Here, we find that Nsp10 activates Nsp16 by opening its SAM and RNA binding loops, allowing them to accommodate their respective ligands. Guided by this activation mechanism, we identify structural states of Nsp16 that are incompatible with substrate binding and also contain potential drug binding sites. Specifically, we find a pocket formed between β3 and β4 of Nsp16 that collapses the SAM binding pocket when open. The region of the pocket has overlap with where Nsp10 binds to Nsp16, so targeting this cryptic pocket could inhibit both substrate (SAM) and Nsp10 binding. Therefore, this cryptic site is a promising target for small-molecule inhibitor development. Further, we find that this cryptic pocket is conserved in MERS and SARS-CoV1 Nsp16, but not in the human homolog CMTr1, suggesting its potential for development of a pan-coronavirus, broad-spectrum inhibitor that may be efficacious against COVID19 and yet unseen coronavirus outbreaks.

## Methods

### System Preparation

The systems were prepared starting from crystal structures 6w4h, 3r24, 5ynf and 4n49, for SARS-CoV-2, SARS-CoV-1, MERS, and CMTr1, respectively. All ligands, solutes, and water molecules from the crystal structures were removed. For monomeric Nsp16 simulations, Nsp10 was also removed. In the coronavirus homologs, two zinc ions were retained, and the coordinating residues were modified accordingly (CYS->CYM and HIS->HID). Missing residues in the crystal structure of CMTr1 were modeled using the Modeller package (39). All systems were solvated in TIP3P water (40) in a rhombic dodecahedral box with periodic boundary conditions and Na^+^and Cl^-^ions added to neutralize the system. Then, systems were energy minimized with a steepest descent algorithm until the maximum force fell below 100 kJ/mol/nm using a step size of 0.01 nm and a cutoff distance of 1.2 nm for the neighbor list, Coulomb interactions, and van der Waals interactions.

Systems were equilibrated for 1.0 ns in *NPT* simulations, with all bonds constrained using the LINCS algorithm (41) and virtual sites were used to allow a 4 s time step. Cutoffs of 1.1 nm were used for the neighbor list with 0.9 for Coulomb and van der Waals interactions. The particle mesh Ewald method (42) was employed for treatment of long-range interactions with a Fourier spacing of 0.12 nm. The Verlet cutoff scheme was used for the neighbor list. Berendsen barostat was used to control the pressure during the equilibration.(43) The stochastic velocity rescaling (v-rescale) thermostat was used to control the temperature at 300 K (44).

### Adaptive sampling simulations

The FAST algorithm (35, 45) was employed for all four homologs for a total of five FAST simulations (SARS-CoV-2 FAST simulations were performed on both monomeric Nsp16 and the Nsp10/Nsp16 complex). FAST was used here to generally enhance conformational sampling and also to quickly explore cryptic pockets. The procedure for FAST simulations is as follows: 1) run initial simulations, 2) build MSM, 3) rank states based on FAST ranking, 4) restart simulations from the top ranked states, 5) repeat steps 2-4 until ranking is optimized. For each system, MSMs were generated after each round of sampling using a k-centers clustering algorithm based on the RMSD between select atoms. Clustering continued until the maximum distance of a frame to a cluster center fell within a predefined cutoff. In addition to the FAST ranking, a similarity penalty was added to promote conformational diversity in starting structures, as has been described previously (46).

For SARS-CoV-2 monomeric Nsp16 and Nsp16/Nsp10, the simulation data was generated in a previous manuscript published by our group. Briefly, FAST-pocket simulations were run at 300 K for 6 rounds, with 10 simulations per round, where each simulation was 40 ns in length (2.4 μs aggregate simulation for each system). The FAST-pocket ranking function favored restarting simulations from states with large pocket openings. Pocket volumes were calculated using the LIGSITE algorithm (47). From these simulations, a conformationally diverse set of structures was selected to be run on Folding@home based on the k-centers clustering algorithm mentioned above. A total of 283 microseconds and 770 microseconds of aggregate simulation time was collected for the Nsp10/Nsp16 complex and monomeric Nsp16, respectively.

FAST-distance simulations were used for SARS-CoV-1 Nsp16, MERS Nsp16, and CMTr1 to sample the β3-β4 pocket identified from SARS-CoV-2 simulations. FAST-distance simulations were run at 300 K for 15 rounds, with 10 simulations per round, where each simulation was 40 ns in length (6.0 μs aggregate simulation for each system). The FAST-distance ranking favored stated with greater distances between the alpha carbons of β3 and β4.

### DiffNets

We used DiffNets, a deep learning-based dimensionality reduction algorithm developed by our group, to highlight biochemically relevant differences between datasets. (36) We trained a DiffNet to compare and contrast structure ensembles of monomeric Nsp16 and the Nsp16/Nsp10 complex to find features that discriminate them, highlighting the structural determinants of Nsp16 activation. First, we subsampled the data by a factor of 25 and 68 for the Nsp16/Nsp10 complex and monomeric Nsp16 data, respectively to have an equal amount of data. Then, we converted simulation data to DiffNet input following the data normalization procedure from the original manuscript. Briefly, XYZ atom coordinates from simulations were mean-shifted to zero, and then multiplied by the inverse of the square root of a covariance matrix, which was calculated from simulations. For all DiffNet training and analysis, we used a split architecture (as described previously) where the classification task was focused on all atoms within 1nm of SAM or RNA-cap based on 6wks crystal structure. This atom selection was chosen to guide DiffNets to find differences in the active site region of Nsp16, which is inherently linked to its activation. For training, simulation frames are classified as “Nsp16 inactive” or “Nsp16 active” based on initial classification labels of 0 (i.e. Nsp16 inactive) for all monomeric Nsp16 frames, and labels of 1 (i.e. Nsp16 active) for all frames from the Nsp10/Nsp16 complex. These labels were iteratively updated in a self-supervised manner described in the original manuscript where we choose expectation maximization bounds of [0.1-0.4] for monomeric Nsp16 and [0.6-0.9] for the Nsp10/Nsp16 complex. This allows for more coherent classification labels as monomeric Nsp16 may sometimes adopt structural poses associated with Nsp16 activation and vice-versa for the Nsp10-Nsp16 complex. Additionally, we used 30 latent variables, 10 training epochs where we subsampled the data by a factor of 10 in each epoch, a batch size of 32, and a learning rate of 0.0001.

To analyze the DiffNet output, we calculated 10 representative structures that span from “Nsp16 inactive” states to “Nsp16 active” states (i.e. structures with classification labels spanning 0 to 1). After training, the DiffNet learns a low-dimensional representation of each simulation frame (i.e. a latent vector) and outputs a classification label for every simulation frame. We binned the structures into 10 equally spaced bins based on their classification labels, which span from 0-1. The, we calculated the mean latent vector for each bin and used the DiffNet to reconstruct a structure based on each latent vector. These structures were used as representative structures for each bin. All training and analysis were performed using the open-source package https://github.com/bowman-lab/diffnets.

### Markov State Models

A Markov State Model (MSM) is a statistical framework for analyzing molecular dynamics simulations that provides a network representation of a free energy landscape. (37, 48, 49) To quantify cryptic pocket opening across the homologs and changes between monomeric Nsp16 and the Nsp10/Nsp16 complex, we performed several measurements that rely on MSMs built based on the simulation data. We built a separate MSM for each system using all simulation data available for that system. All MSMs were constructed with the Enspara python package (50). First, the solvent accessible surface area (SASA) of each residue side-chain was calculated using the Shrake-Rupley algorithm (51) implemented in MDTraj (52) using a drug-sized probe (2.8 Å sphere).

Then, we clustered the data using a hybrid clustering algorithm. First, we used a k-centers algorithm (53) to cluster the data (5000 clusters for SARS-CoV-2 data, 1500 clusters for each other homolog). Next, we applied sweeps of k-medoids update steps (3 for SARS-CoV-2 data, 2 for other homologs) which refined the cluster centers to be in the densest regions of conformational space (54). A Markov time of 5 ns was selected for based on the implied timescales to build a Markov state model (MSM) for each homolog. To build the MSMs, transition probability matrices were produced by counting transitions between states (i.e. clusters), adding a prior count of 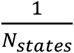 and row-normalizing, as is described previously (55). Equilibrium populations were calculated as the eigenvector of the transition probability matrix with an eigenvalue of one. For all histograms shown, we calculated the order parameter of distance (e.g. distance between β3-β4) using cluster centers (i.e. representative structure of the cluster) and weighted the order parameter by the corresponding equilibrium population calculated with the MSM. We also resampled the equilibrium populations 100 times by bootstrapping the MSM, which provided error bars for computing the fraction of SAM and RNA compatible states adopted by monomeric Nsp16 and the Nsp16/10 complex.

### Distance and SASA calculations

Figures 2, 3, and 4 include distance and SASA measurements that are explained in more detail here. In Figure 2 we measure the distance between gate loop 1 and gate loop 2 as the distance between Gln28 and Lys141 since these residues are known to undergo significant changes for RNA binding. We measure the distance between SAM binding loop 2 and gate loop 2 as the average distance between (Met131, Tyr132, Asp133, Pro134) and (Asp99, Leu100, Asn101, Asp102) as these are key residues that cradle SAM in the bound state. All SASA measurements are performed using Ala79, Thr82, Ala83, Leu86, Thr93, Leu94, Leu95, Val96, Asp97, Ala98 and Asp99 as this is the main component that gets exposed during cryptic pocket opening.

### Cryptic pocket detection

Cryptic pockets in SARS-CoV2 Nsp16 were identified using our previously established approach called Exposons analysis (38). This analysis was performed using the cluster centers and the equilibrium probabilities derived from the MSMs built on the residue level SASA described above. The center of each cluster was taken as an exemplar of that conformational state, and residues were classified as exposed if their SASA exceeded 2.0 Å^2^ and buried otherwise. The mutual information between the exposure/burial of each residue-pair was calculated based on the MSM, by treating the SASA values in the cluster centers as samples and weighting them by the equilibrium probability of the representative state. The mutual information was computed using the following equation:

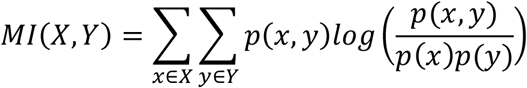

Finally, cryptic pockets (Exposons) were identified as groups of residues undergoing cooperative change in SASA, by clustering the matrix of pairwise mutual information using affinity propagation.

The β3-β4 cryptic pocket identified in SARS-CoV2 Nsp16 consists of residues Ala79, Thr82, Ala83, Leu86, Thr93, Leu94, Leu95, Val96, Asp97, Ala98 and Asp99. Total SASA of these residues/homologous residues was measured for detecting cryptic pocket opening in all homologs of Nsp16 (SARS-CoV2, SARS-CoV1 and MERS). For measuring equivalent cryptic pocket in CMTR1, total SASA of structurally homologous residues (Gly141, Ser144, Glu145, Val148, Ala155, Lys156, Gly157, His158, Gly159, Met160, Thr161) was calculated.

### Sequence Conservation

Protein sequences of Nsp16 from SARS-CoV2 (YP_009725311.1), SARS-CoV1 (Uniprot ID: P0C6×7), MERS (Uniprot ID: K0BWD0), NL63 (AFD64750.1), HKU1 (YP_460023.1), Turkey CoV (YP_001941189.1), Bat CoV (YP_008439226.1), Murine hepatitis virus (YP_209243.1) were used for multiple sequence alignment. Sequences alignment was performed on Clustal Omega server (56). Sequence alignment was visualized, and the sequence conservation score was generated using Jalview 2 software (57).

For sequence comparison of SARS-CoV2, SARS-CoV1, MERS and human CMTr1 shown in Fig. 4, structure-based sequence alignment was performed using UCSF Chimera package (58). For the structure-based sequence alignment, we first aligned the structures of these homologs (PDB: 6wks (SARS-CoV2), 3r24 (SARS-CoV2), 5ynf (MERS) and 4n49 (CMTr1). Then, the sequences were aligned based on the structural alignment of the backbone atoms.

## Author Contributions

NV, MDW, SS, GRB designed the research. NV, MDW, MIZ, BN, JHB, and SS performed research. NV, MDW, JHB, and GRB wrote the manuscript.

## Acknowledgements

We are grateful to the citizen scientists who contribute to Folding@home by running simulations on their personal computers. This work was funded by NSF CAREER Award MCB-1552471, NSF RAPID 58628, and NIH R01 GM124007. GRB holds a Career Award at the Scientific Interface from the Burroughs Wellcome Fund and a Packard Fellowship for Science and Engineering from The David & Lucile Packard Foundation. We are grateful to Catherine Knoverek for her thoughtful feedback when editing initial drafts of the writing.

## Supporting Information

**Figure S1.**
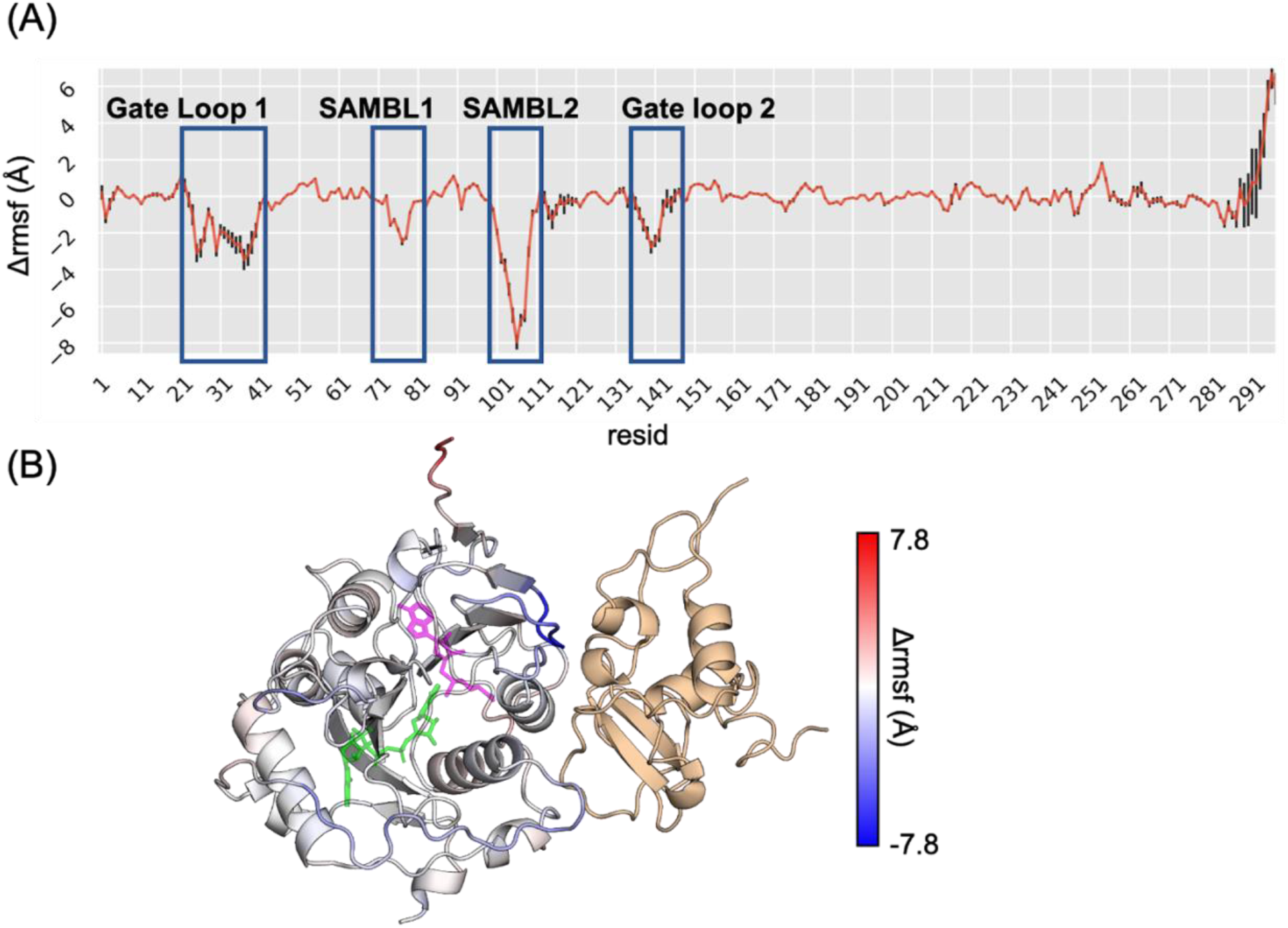
Change in root mean square fluctuation (rmsf) of Nsp16 upon Nsp10 association. (A) Probability weighted Δrmsf of Nsp16’ residues upon Nsp10 binding is plotted. Negative values represent a decrease in rmsf upon Nsp10 binding. RNA binding loops (gate loop 1 and 2) and SAM binding loops (SAMBL1 and 2) are highlighted by the blue colored boxes. (B) Probability weighted Δrmsf of Nsp16 is mapped on its structure, with negative values shown in blue and positive values in red.

**Figure S2.**
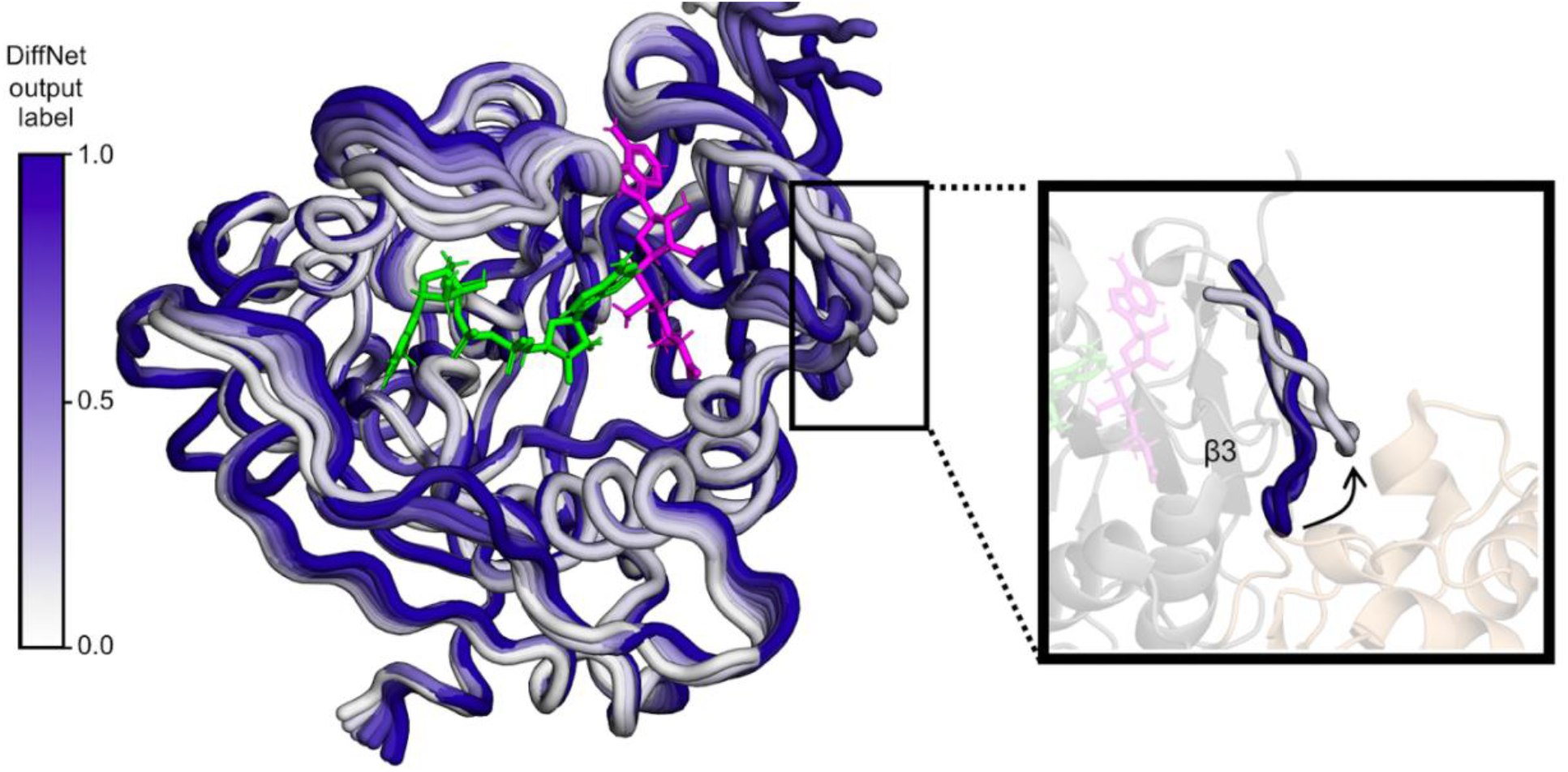
DiffNets predict that β4 peels away from β3 in Nsp16 inactive structural states. (Left) Structural states changing from inactive to active (white to purple) as predicted by the DiffNet. (Right) The loop connecting β3 and β4 peels away from β3 into solution in predicted inactive states.

**Figure S3.**
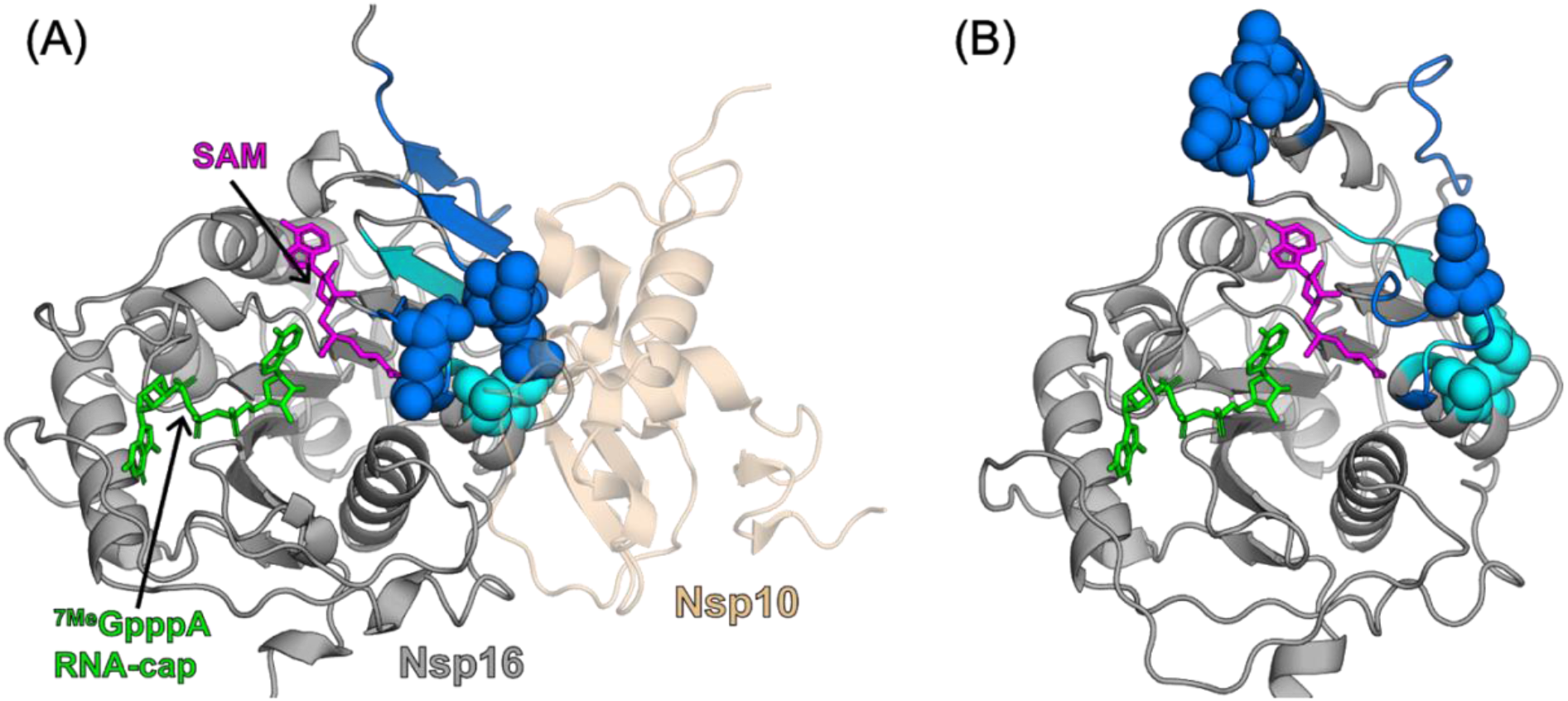
Displacement of Nsp10 binding residues by cryptic pocket opening. (A) Structure of Nsp16 in cryptic pocket closed state is shown in grey. Cryptic pocket forming residues and the residues undergoing opening motion are shown in cyan and blue, respectively. Cryptic pocket residues that contact Nsp10 are depicted in spheres. (B) Opening motion of the cryptic pocket shows the displacement of Nsp10 binding residues.

**Figure S4.**
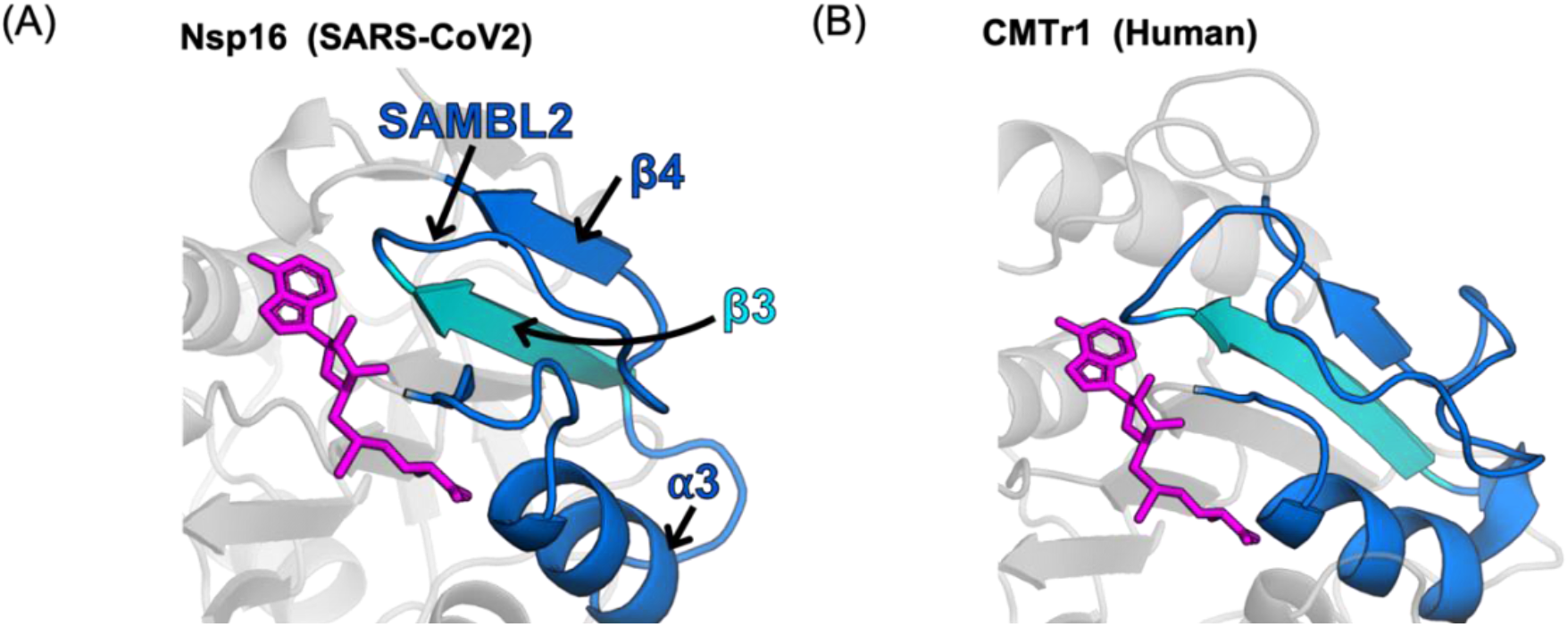
Structural comparison of β3-β4 cryptic pocket in SARS-CoV2 Nsp16 and human CMTr1. (A) β3 and residues lining the cryptic pocket in SARS-CoV2 are shown in cyan and blue, respectively. (B) Regions of human CMTr1, structurally equivalent to β3 and the pocket lining regions are depicted in cyan and blue, respectively.

**Figure S5.**
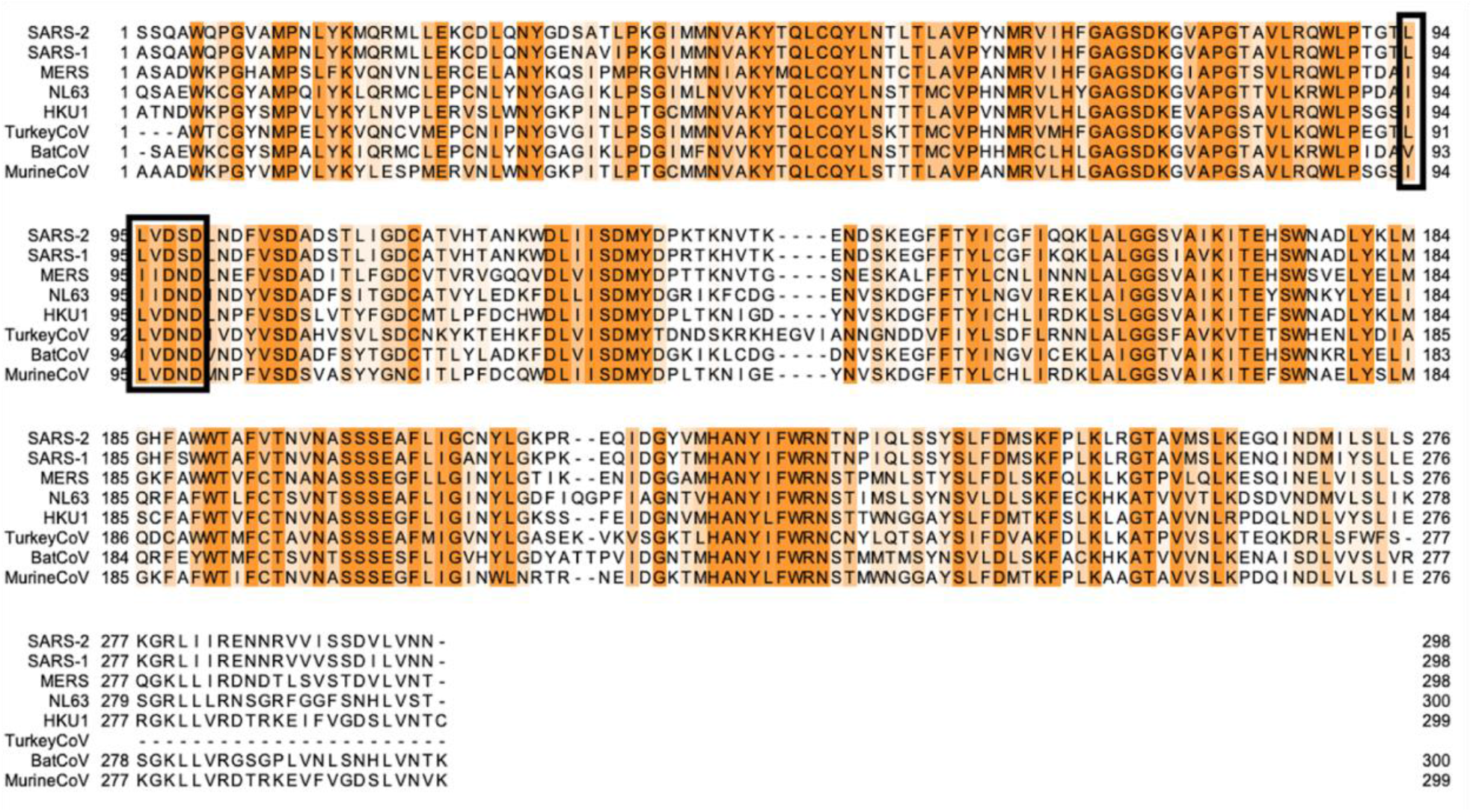
Multiple sequence alignment of Nsp16 homologs from coronaviruses. The color ranges from white to orange for the sequence conservation score ranging from 0 to 10, where 10 denotes 100% sequence identity. Residues of β3 are enclosed in the black box. Uniprot ids of the sequences used for the alignment are given in the Methods section.

